# Transcriptome Analysis of NPFR Neurons Reveals a Connection Between proteome Diversity and Social Behavior

**DOI:** 10.1101/2020.11.12.379826

**Authors:** Julia Ryvkin, Assa Bentzur, Anat Shmueli, Miriam Tannenbaum, Omri Shallom, Shiran Dokarker, Mali Levi, Galit Shohat-Ophir

## Abstract

Complex social behaviors are mediated by the activity of highly intricate neuronal networks, the function of which is shaped by their transcriptomic and proteomic content. Contemporary advances in neurogenetics, genomics, and tools for automated behavior analysis make it possible to functionally connect the transcriptome profile of candidate neurons to their role in regulating behavior. In this study we used *Drosophila melanogaster* to explore the molecular signature of neurons expressing receptor for neuropeptide F (NPF), the fly homologue of neuropeptide Y (NPY). By comparing the transcription profile of NPFR neurons to those of nine other populations of neurons, we discovered that NPFR neurons exhibit a unique transcriptome, enriched with receptors for various neuropeptides and neuromodulators, as well as with genes known to regulate behavioral processes, such as learning and memory. By manipulating RNA editing and protein ubiquitination programs specifically in NPFR neurons, we demonstrate that their delicate transcriptome and proteome repertoires are required to suppress male courtship and certain features of social group interaction. Our results highlight the importance of transcriptome and proteome diversity in the regulation of complex behaviors and pave the path for future dissection of the spatiotemporal regulation of genes within highly complex tissues, such as the brain.

## Introduction

Behavior is the result of an orchestrated neuronal activity, where a complex repertoire of cell types assembled into circuits processes external and internal information into a consistent motor output that ultimately promotes survival and reproduction^1–4^. The immense complexity and heterogeneity of the nervous system results from molecular programs that dictate the repertoire of expressed proteins, including their localization and function, giving rise to cell populations with diverse anatomy, physiology, connectivity, and functional roles^5–13^. This diversity poses a challenge when trying to functionally associate neurons to particular behaviors but can be resolved by genetically dividing the brain into discrete cell types and subsequently study their anatomy, connectivity, molecular architecture and physiology^14–18^. Recent advances in targeting increasingly smaller sub populations of neurons, together with tools to manipulate their activity, make it possible to connect the function of neurons to their identity, thus facilitating greater understanding of the molecular underpinning of brain development and mechanisms that regulate complex behaviors^19–23^. This can be useful when studying the function of neurons that control complex behaviors, particularly those that are regulated by motivation such as foraging, food and water consumption, mating and various forms of social interactions^2,24–30^.

The fruit fly *Drosophila melanogaster* is a useful model organism for investigating the genetic underpinnings of motivational behaviors, owing to variety of tools for neuro-genetic manipulations, together with the fact that flies exhibit several forms of behaviors that are shaped by motivation^29,31–41^. One of the systems that encodes internal states and dictates motivational drives, and consequently, behavioral choices in *Drosophila* is the Neuropeptide F/Neuropeptide F receptor^42–54^. Most studies in the field have focused on NPF-producing neurons and less on NPFR-expressing neurons. Similar to their mode of action in mammalian systems, NPFR neurons are inhibited by NPF^48,55–58^ and are positioned in a convergence point for the regulation of various complex behaviors including male sexual behavior^45,46^, ethanol consumption and sensitivity^42,51,59^, feeding^44,54^, appetitive memory^33^, arousal and sleep^60,61^. Nevertheless, the molecular basis underlying their diverse function is largely known.

In this work, we investigated the transcriptional landscape of NPFR neurons, comparing it to those of nine other neuronal populations, and discovered that NPFR neurons have a unique signature that is enriched in neuropeptide and neuromodulator receptors. We tested the functional relevance of this intricate transcriptome and proteome repertoire by disturbing RNA editing and protein ubiquitination programs in NPFR neurons. Our results show that these manipulations enhance certain aspects of male-female and male-male interactions, suggesting a role for NPFR neurons in restraining social and sexual behaviors.

## Results

To explore the connection between transcriptional programs in NPFR neurons and behavior, we used a recently generated dataset from our lab consisting of RNA sequences from several neuronal populations in the brain, that were obtained by immunoprecipitation of genetically tagged nuclei (INTACT method)^8^. The dataset included 10 neuronal populations known to regulate behavior and physiology: dopaminergic neurons (*TH-Gal4*), octopaminergic neurons (*Tdc2-Gal4*), serotonergic neurons (*TRH-Gal4*), NPF neurons (*NPF-Gal4*), NPF-receptor neurons (*NPFR-Gal4*), mushroom bodies (*OK107-Gal4*), Corazonin neurons (*CRZ-Gal4*), Dh44 neurons (*CRF* orthologue *DH44-Gal4*), fruitless-expressing neurons (*Fru-Gal4*), and all neurons (*Elav-Gal4*). Analysis of transcriptomic datasets offers a way to compare the levels of transcription per gene across different cell populations, or within the same cells under different conditions. To explore the transcriptomic landscape of NPFR cells, we took two complementary approaches: pairwise comparison of gene expression profiles between each neuronal population and all neurons (pan-neuronal driver); and pairwise comparison of gene expression profiles between each neuronal population and NPFR neurons.

### The Transcriptomes of NPFR, Fru, and OK107 Neurons are Most Similar to Those of the General Neuronal Population

Starting with the first approach, we generated a list of genes for each neuronal population with significantly different expressions than those in all neurons (greater than two-fold change compared to the expression in ElaV), called “yes genes” (Figure 1A, B; Table S1). Since the amount of yes genes in each neuronal population represents the difference in transcriptome between this population and all neurons, we expected that the more specific the transcriptome in a population is, the more unique it will be compared to ElaV. Interestingly, DH44- and NPF-expressing neurons displayed the largest number of yes genes (2758 and 1990 respectively), while OK107- and NPFR-expressing neurons presented the smallest number of yes genes (40 and 42, respectively) (Figure 1A). Most yes genes in OK107, NPFR, TRH, Tdc2, and TH were found to be enriched compared to those in ElaV, while most yes genes in Fru neurons were depleted compared to those in ElaV (Figure 1A). Hierarchical clustering analysis of average normalized reads for all the yes genes between the different neuronal populations confirmed this finding: DH44 cells were clustered apart from all other populations, followed by NPF cells (Figure 1B); in addition, OK107 cells clustered closest to ElaV, together with Fru and NPFR neurons (Figure 1B). Altogether, this suggests that the transcriptomes of DH44- and NPF-expressing cells are the most unique, while those of OK107-, Fru-, and NPFR-expressing neurons are similar to the transcriptome in the general neuronal population.

**Figure 1:**
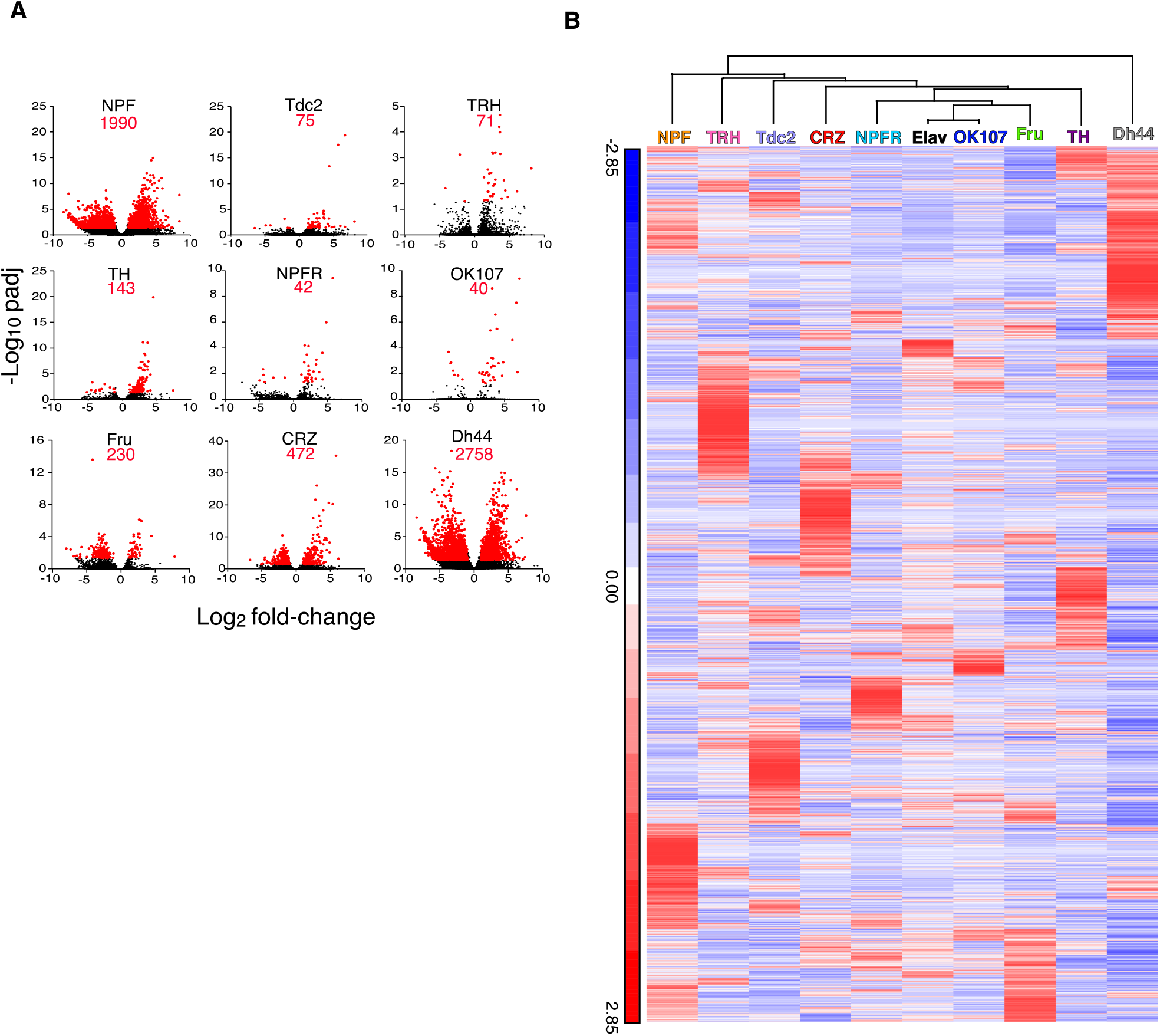
Different neuronal populations exhibit varying number of differentially expressed genes when compared to the general population of neurons. **A** Volcano plots of log_2_ average fold change per population of all genes (black) and significantly expressed genes (red) compared to ElaV. **B** Hierarchical clustering of average normalized reads for all significantly expressed genes in 9 neuronal populations compared to a pan neuronal driver (ElaV). Clustering analysis was performed using Partek.

### Shared Yes Genes between Populations Reveal a Complex Pattern

Given the partial anatomical overlap between several neuronal populations in our dataset^46,15,62–65^, we next asked whether some yes genes are shared across different neuronal populations. Enrichment or depletion of the same genes in more than one population suggests that these neuronal populations share differences from the general population, and/or that some of their neurons overlap. Searching for yes genes that are shared by different neuronal populations, we did not document any genes that are shared by all nine populations (Figure 2A, Table1, TableS2). When comparing shared yes genes across 8-3 neuronal populations, only a single gene (CG9466) was found to be shared by eight populations, exhibiting similar pattern of enrichment in all eight populations (Figure 2A and Table S2). The long non-coding RNA CR45456 is another example for a transcript that is enriched in six neuronal populations when compared to its expression ElaV (Figure 2A, Table S2).

**Figure 2:**
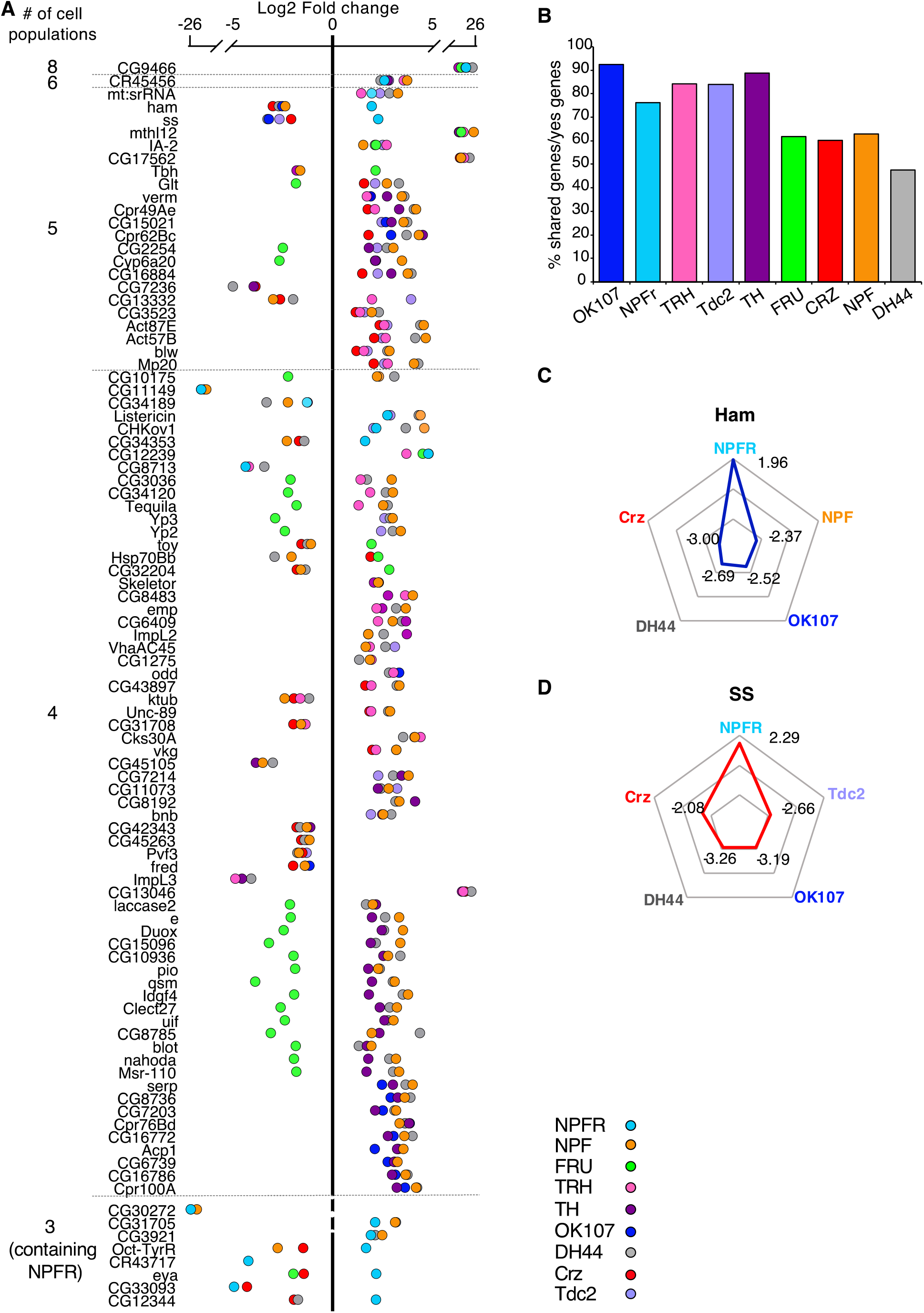
Certain yes genes are shared across many populations. **A** Scatter plot representing log_2_-fold change of all genes that are differentially expressed compared to ElaV (yes genes) and are shared across 8-4 different cell populations (upper part) and across 3 populations containing NPFR (bottom part). **B** Percent of shared yes genes normalized by total number of yes genes in each neuronal population. **C,D** Radar plots of two yes genes: ham(C) and ss(D) both of which were enriched in NPFR cells compared to 4 other cell types.

**Table 1:**
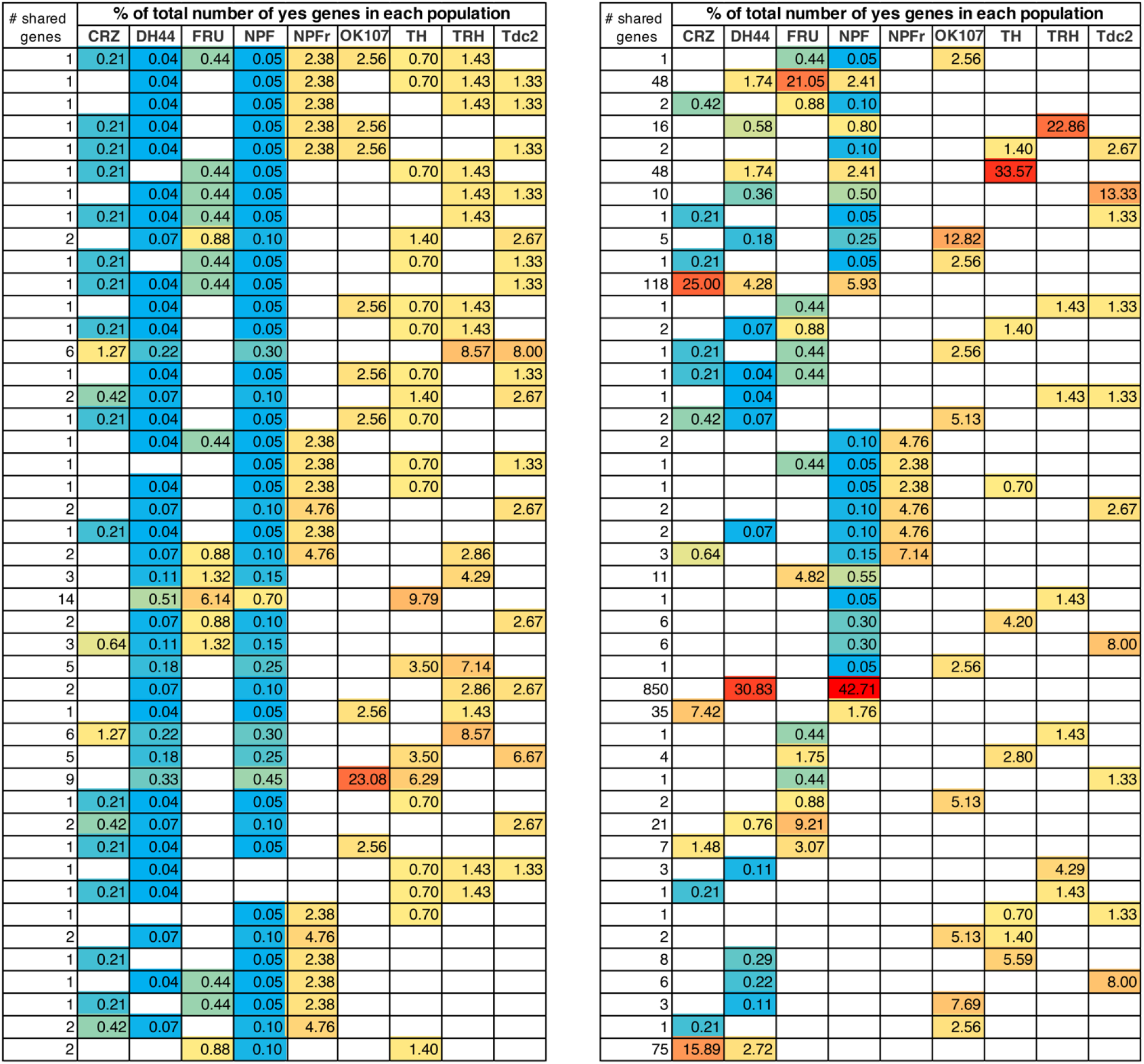
The amount of shared yes genes varies across populations. Number of shared yes genes across populations (left column) represented as % from the total amount of yes genes in each population (color coded: red-high, blue-low).

Two neuronal populations shared the largest number of yes gene with all other populations; NPF and DH44 (1253 genes in 67 comparisons and 1309 genes in 56 comparisons, respectively, Figure 2A, Table 1, Table S2). The amounts of yes genes varied across all populations by two orders of magnitude (Figure 1A), increasing the odds for shared yes genes in certain populations due to the overall number of yes genes and not because they were expressed within overlapping neurons. To control for this, we normalized the amount of shared yes genes by the total number of yes genes in each population and found a reduction in the variation of the numbers of shared yes genes between populations (Figure 2B). This finding implies that the probability of sharing a yes gene is similar across different populations, and that the more yes genes a population has, the higher the probability that some will be shared, emphasizing the need to use other criteria to determine whether two populations share similar transcriptional patterns or just mutual neurons.

Interestingly, and although Fru shares neurons with several other populations, such as NPF and Tdc2^46,63,65^, as evidenced by the enrichment of Tbh (Tyramine β hydroxylase) in Fru and Tdc2 neurons, most yes genes in Fru neurons were depleted compared to their expressions in other populations (Figure 2A). Striking examples are Cyp6a20, Glutactin, Tequila, toy, and quasimodo, which support the notion that most Fru neurons are distinct from the rest of the analyzed neuronal populations. In addition, CRZ-, DH44- and NPF-expressing neurons shared similar expression patterns of groups of genes that shape neurophysiology, possibly due to all of them being peptidergic neurons. Examples of these neurophysiology-associated genes include: the shared patterns of ion channels, such as NaCP6OE (Voltage gated Na channel), Teh1 (TipE homologue 1 sodium transport regulation), genes involved in neuronal signaling, such as Neuroligin 3 (synaptic adhesion molecule), beat-1C (beaten path 1C axon guidance), Tehao (Toll signaling); and the shared patterns of receptors, such as nicotinic acetylcholine receptor alpha3 and 6, Toll6 (Toll-like receptor family), IR47a+b (ionotropic receptor a+b), GluR1A (Glutamate receptor 1A), and Oct-beta-3R (Octopamine receptor beta 3).

The shared yes genes between NPFR neurons and other neuronal populations illuminated a complex pattern of 21 genes that are similarly and oppositely expressed (Figure 2A). The two most differentially regulated genes were hamlet (ham) and spineless (ss), both highly enriched in NPFR neurons and depleted in all other neuronal populations (Figure 2C, D). Examining shared yes genes in comparison to NPF neurons revealed two more genes with opposite expression that are enriched in NPFR (Octopamine-Tyramine Receptor and CG34353) and 11 yes genes with similar expression. NPFR neurons also displayed expression patterns of yes genes different from those in DH44 neurons, with four oppositely expressed yes genes, including ham, ss, CG34353 and CG12344, and similarly expressed genes, like CG9466, CR45456, mt:srRNA, CG10175, CG34189, Listericin, CHKOV1, CG12239, CG8713, CG31705, CG3921, CR43717 and CG33093 (Figure 2A). Interestingly, Octopamine-Tyramine Receptor (Oct-TyrR), which is regulated by feeding and mediates appetitive changes in locomotion^66^, was enriched in NPFR-expressing neurons and depleted in NPF- and CRZ-expressing neurons (Figure 2A). Furthermore, ss, which encodes a transcription factor regulating female receptivity to male courtship^67^, was enriched in NPFR neurons. This data suggests that while it is possible that some of the NPF and DH44 neurons share neuronal subpopulations with NPFR, many of the neurons in these populations do not overlap.

Next, we analyzed the relative expression patterns of NPFR neurons using the second pairwise approach; comparing NPFR neurons to each of the neuronal populations (Figure 3, Table S3). The pairwise comparison of NPFR to ElaV expression profiles resulted in the identification of 42 yes genes, but comparing the expression pattern of NPFR neurons to those of all other populations revealed a larger number of differentially-expressed genes, with up to 2669 differentially-expressed genes between NPFR and DH44 neurons. Interestingly, with the exception of the DH44 neurons, all other cell populations presented higher amounts of differentially-expressed genes when compared to NPFR neurons than when compared to ElaV neurons. This could suggest that DH44 neurons are more similar to NPFR neurons than to the general population neurons (Table S3). Hierarchical clustering analysis for the identified yes genes in each of the comparisons revealed that DH44- and NPF-expressing neurons clustered away from the rest of the populations, while Fru and OK107 neurons were most similar to NPFR neurons (Figure 3), with a clustering pattern similar to that obtained when comparing them to ElaV in Figure 1B.

**Figure 3:**
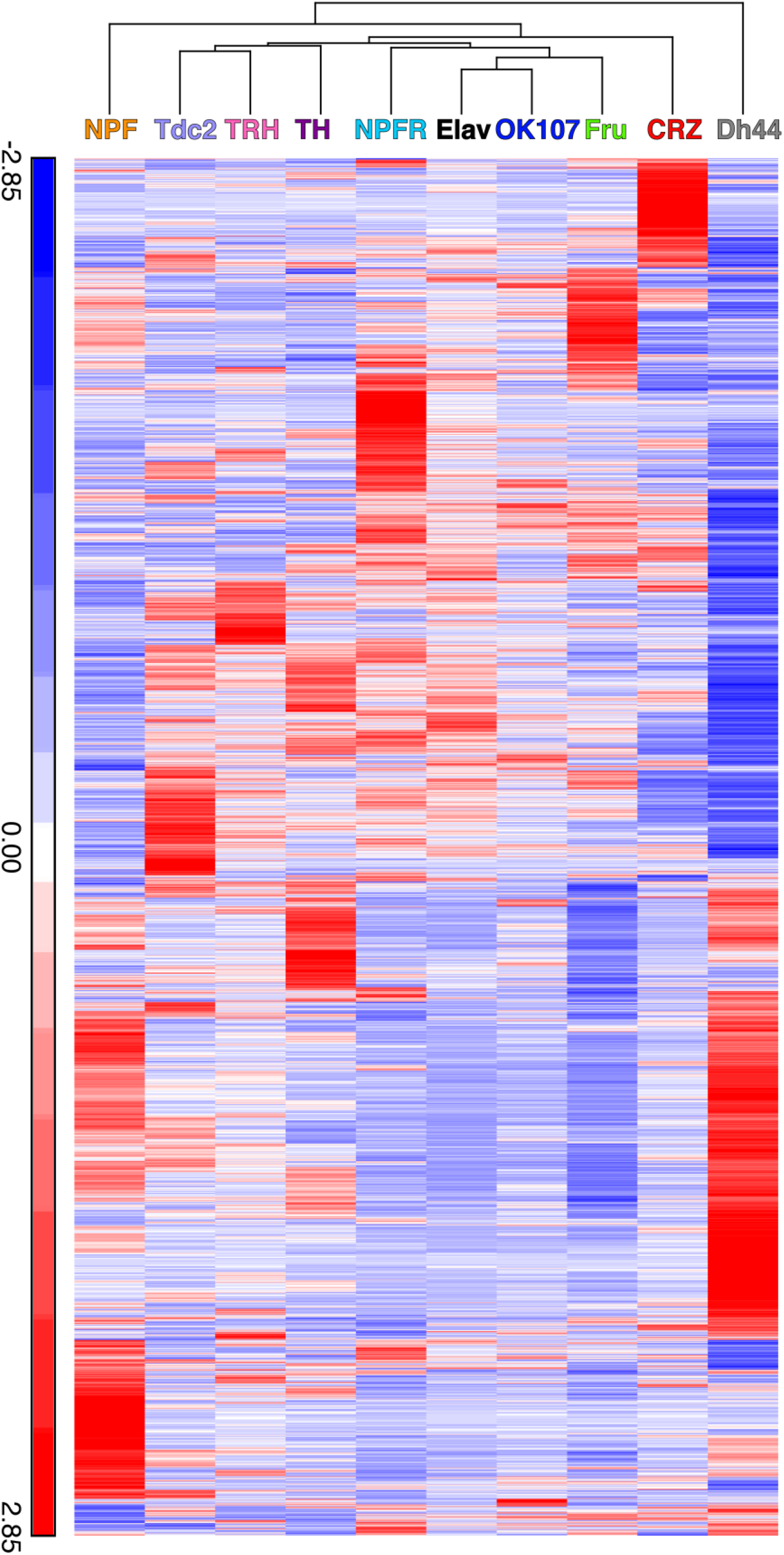
Hierarchical clustering of average reads for all differentially expressed genes compared to the NPFR neurons. Clustering analysis was performed using Partek.

To further explore the biological relevance of the identified yes genes, we used a statistical overrepresentation analysis (PANTHER), which highlighted several biological processes, including enrichment of genes associated with regulation of behavior (Figure 4, Table S4). We focused on behavior-associated genes that were enriched or depleted in NPFR vs. CRZ, TH, Fru, and OK107 neurons and found some interesting patterns (Figure 4). NPFR neurons displayed enrichment of genes that mediate different forms of learning and memory, such as derailed, 2mit, klingon, CG18769, Omab, mGluR (metabotropic Glutamate Receptor), eag, Ank2, ss, and Tequila (Figure 4). In addition, we identified enrichment of genes involved in sensory perception of sound and touch, such as Ank2, btv, nompC, CG14509, DCX-EMAP, dila, and Rootletin. Interestingly, we documented enrichment of a few genes that participate in insulin signaling, such as dilps 2, 3, and 5 in Dh44 neurons, suggesting an anatomical overlap between some NPFR neurons and insulin-producing cells (IPCs).

**Figure 4.**
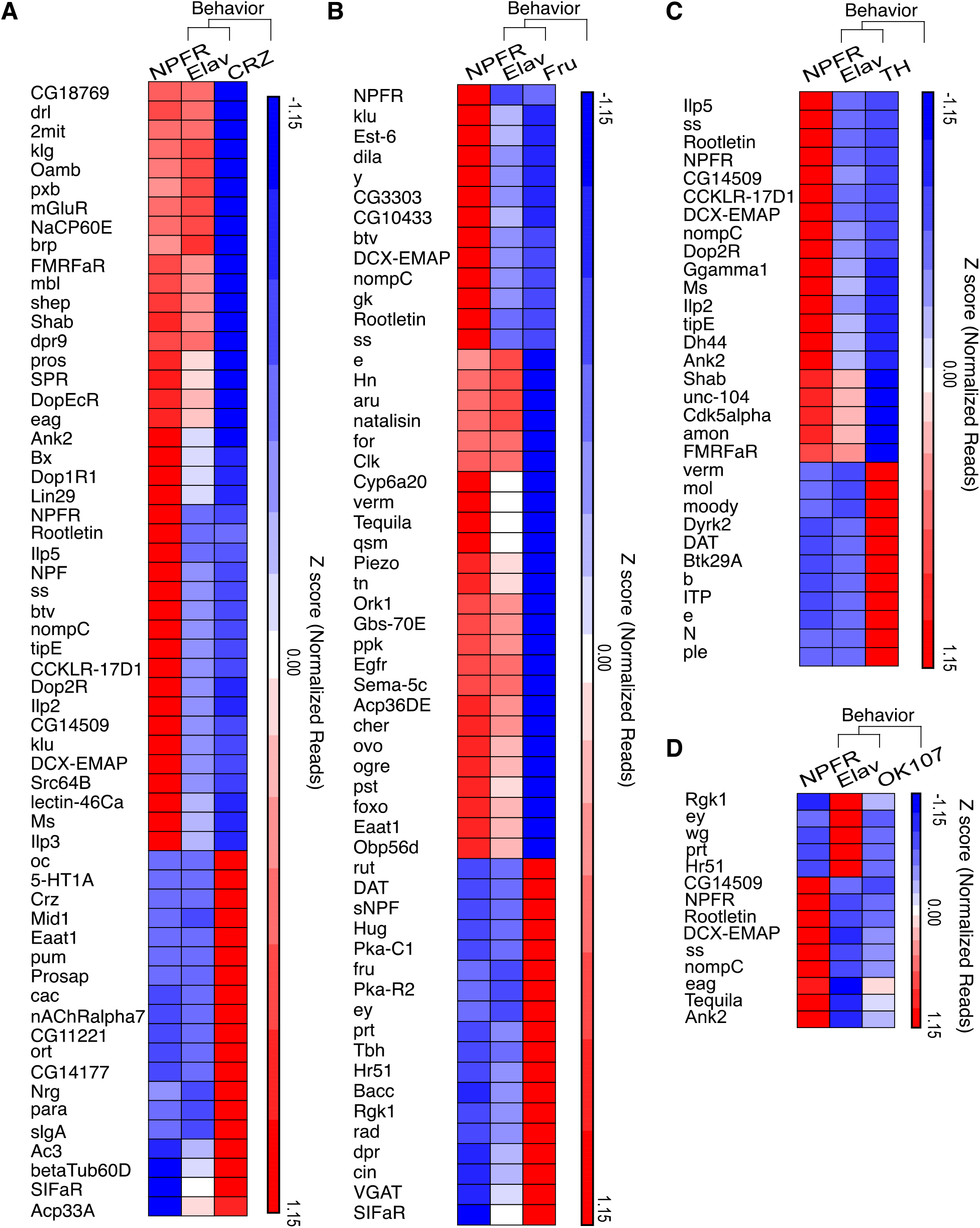
NPFR expressing neurons exhibit enrichment of behavior related genes. **A-D** Hierarchical clustering of statistically overrepresented biological processes that are related to behavior in differentially expressed genes of NPFR expressing neurons compared to Corazonin (CRZ, **A**), Fruitless (FRU, **B**), Dopamine producing (TH, **C**) neurons and Mushroom bodies neurons (OK107, **D**). Biological overrepresentation was performed using PANTHER. Clustering analysis was performed using Partek.

Intriguingly, NPFR neurons exhibited enriched levels of various receptors for neuropeptides and neuromodulators like Oamb, mGluR, Dop1R1, Dop2R, CCKLR-17D1, MS, Lestin-46Ca, Ms, TrissinR, CCHa1-R (CCHamide-1 receptor), AstA-R1(Allatostatin A receptor1), rk (rickets), Proc-R (Proctolin receptor), SPR (sex peptide receptor), sNPF-R, and (of course) the receptor for NPF (Figure 4, S1). The enrichment of such a diverse repertoire of receptors indicates that NPFR neurons receive multiple inputs from many neuromodulator systems, and/or that they are composed of diverse groups of neurons, with distinct combinations of receptors. In any event, these findings support the hypothesis that NPFR neurons are located at a convergence point of information that is relevant for the integration of internal state and action selection.

Lastly, we found that NPFR neurons possessed a unique mixture of ion channels compared to both Crz and Fru neurons (Figure S3), with an overrepresentation of seven potassium and sodium ion transmembrane transport subgroups in NPFR neurons (Table S4). We also documenter an overrepresentation of amino acid transmembrane transport proteins with 2 enriched genes in Fru cells (vGAt, CG5549) and 6 enriched in NPFR neurons (Ncc69, CG7888, Eaat1, CG43693, CG8785, CG16700). Interestingly, Orct2 (Organic cation transporter 2), which is a transcriptional target of the insulin receptor pathway^68^ (Figure S2) was enriched in NPFR compared to Crz, further supporting the involvement of NPFR neurons in insulin signaling.

### Manipulation of the Proteome Profile in NPFR Neurons Affects Social and Sexual Behavior in Flies

The distinct patterns of transcription in each neuronal population gives rise to a specific proteome diversity that shapes the functional output of neurons. To investigate this assumption further, one can modify the expression levels of genes that are enriched or depleted in certain populations or use a more global approach to disturb the delicate proteomic signature of the neurons. We chose to perturb the transcriptomic and proteomic signature of NPFR neurons by manipulating the function of two molecular systems that regulate a large number of cellular targets (RNA editing and protein ubiquitination) and to analyze the effects on the social behavior of male flies.

Adenosine-to-inosine (A-to-I) RNA editing, catalyzed by ADAR enzymes^69,70^, is a cellular mechanism that generates transcriptomic and proteomic diversity by recoding certain adenosines within pre-mRNA sequences into inosines, leading to a variety of consequences that include amino acid sequence changes in proteins. Thousands of RNA editing sites have been discovered in *Drosophila*^71^, most lead to recoding events in genes that are expressed and function specifically in the neuron^71–76^. As such, null mutation of ADAR in *Drosophila* leads to strong locomotor phenotypes that become more severe with age, the underlying mechanisms is still mostly unknown^77^. Therefore, we hypothesized that reducing ADAR expression in NPFR neurons would affect the proteomic profile and could result in behavioral phenotypes. To test this, we downregulated the expression of dADAR in NPFR neurons (NPFR>UAS-dicer, UAS-dADAR RNAi) and analyzed behavior in groups of 10 flies, using the “FlyBowl” system, a suite of tracking and behavior analysis softwares that score plethora of locomotion and social behaviors^78,79^. We used the tracking data obtained to generate a comprehensive behavioral representation for experimental flies and genetic controls that included kinetic features and eight complex behaviors. The overall differences between the genotypes are depicted in a scatter plot of normalized differences, divided into four main categories: activity-related features, interaction-related features, coordination between individuals, and social clustering-related features (Figure 5A).

**Figure 5.**
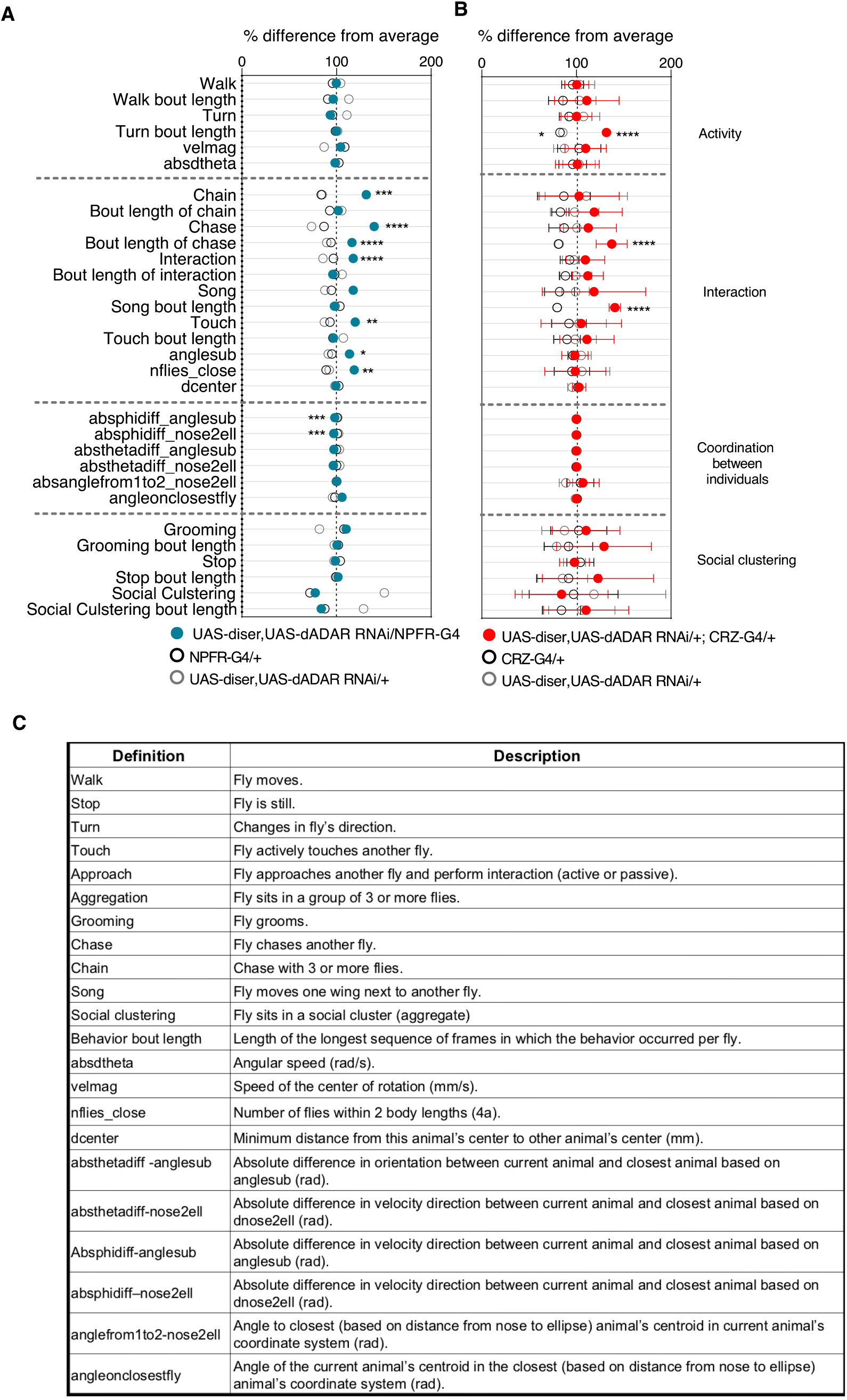
RNA editing is required in NPFR neurons for typical male social behavior. Behavioral signatures of social group interaction. Data is presented as normalized scatter plots depicting % difference from average of 31 behavioral features. **A** behavior profile of male flies harboring NPFR-Gal4/ UAS-Dicer, UAS-dADAR RNAi (Blue) compared to genetic controls (UAS-Dicer, UAS-dADAR RNAi/+, and NPFR-Gal4/+, grey and black respectively). n=17. **B** behavior profile of male flies harboring UAS-Dicer, UAS-dADAR RNAi/+; CRZ-Gal4/+ (red) compared to genetic controls (UAS-Dicer, UAS-dADAR RNAi/+, CRZ-Gal4/+, grey and black respectively). n=7. One-way ANOVA with Tukey’s post hoc for normally distributed parameters and Kruskal-Wallis with Friedman test post hoc for non-normally distributed parameters. *P<0.05, **P<0.01, ***P<0.001, ****p<0.0001 n.s. P>0.05.

Unlike dADAR null flies and pan-neuronal knockdown (KD) of dADAR flies, which display strong motor impairments, downregulation of dADAR expression in NPFR neurons did not lead to any differences in locomotion and general activity levels. Specifically, the average velocity of experimental flies and the percentage of time they spent walking and performing body turns was similar to those of the genetic controls (NPFR-Gal4/+ and UAS-dicer, UAS-dADAR RNAi/+, Figure 5A). We further analyzed several types of social behaviors, including touch (active leg touching between two flies), approach (fly approaching another fly), song (wing extension and vibration to generate male courtship song), chase (fly chasing another fly), and chaining (one fly following a fly while being followed by another fly, in a minimum chain length of three flies). Interestingly, reducing ADAR levels in NPFR neurons resulted in strong elevation in social interaction between male flies, as manifested by increased levels of close touch behavior, increased levels of song displays, increased values of active approaches and male-male chase events that resulted in multiple formations of chains (Figure 5A). In addition to these behaviors, we analyzed two other features associated with social interaction: angle subtended (anglesub), representing the maximum angle of an animal’s field of view occluded by another animal as a measure of the distance between flies; and the number of flies close-by (nflies-close), representing the number of flies within two body lengths of a focal fly as a measure of sociality (Figure 5A,C). Flies harboring reduced levels of ADAR in NPFR neurons depict significantly higher values in these two features compared to the control groups, suggestive of close distance between flies (Figure 5A), altogether indicating that RNA editing in NPFR expressing cells is important for the correct expression of certain social behaviors. A previous study in our lab demonstrated that NPFR and CRZ neurons possess distinct RNA editing repertoires^8^. This prompted us to test the behavioral significance of reducing RNA editing in CRZ neurons as well. However, knocking down ADAR in CRZ neurons only led to moderate effects on the male-male social interactions. Specifically, we documented longer bouts of of song, turn, and chase events than in genetic controls (Figure 5B).

Next, we perturbed the intricate proteome diversity in NPFR cells by targeting the protein ubiquitination machinery, a highly important mechanism that is necessary for the regulation of protein degradation and function. To manipulate this multiplayer system, we targeted the expression of one central player, ubiquitin-conjugating enzyme 7 (Ubc7), orthologous to the human ubiquitin-conjugating enzyme E2 G2. We used the CrispR-Cas9 system to generate tissue specific knockout (KO) of Ubc7 using a combination of Ubc7-specific guide RNAs and specific expression of Cas9 in NPFR neurons (NPFR>UAS Cas9)^80–82^. We generated a pair of guide-RNAs (gRNA) targeting the beginning of the second axon of Ubc7. We validated their efficiency by using a germline deletion within the Ubc7 locus by driving Cas9 expression using a germline-specific driver (Vas-Cas9), which resulted in an 18-to 23-bp deletion at the beginning of the second axon of both Ubc7 isoforms (Figure 6A, B S3). To affect Ubc7 in NPFR cells, we crossed *NPFR*-G4; *UAS-Cas9*.c flies with flies carrying our gRNA for Ubc7. Since Ubc7 null mutation was shown to suppress courtship towards females^83^, we first analyzed the effects of knocking out Ubc7 in NPFR neurons on male courtship behavior. For that, we introduced experimental male flies (NPFR>UAS Cas9, *gRNAs*) or genetic control male flies (*NPFR-G4/attp1; UAS-Cas9.c*) into courtship arenas with virgin females and recorded and analyzed their behavior (Figure 6C-E). Surprisingly, and in contrast with Ubc-7 null mutants, male flies lacking Ubc7 expression in NPFR cells displayed shorter latency to court, shorter latency to first copulation attempt, and shorter time to successful copulation (Figure 6C-E), all signs suggestive of higher motivation to court and mate. Next, we analyzed the behavioral responses of male flies when interacting with nine other male flies in a group (Figure 6F). Contrary to the previous results obtained after manipulating the proteome of NPFR by disturbing RNA editing programs, which did not affect any of the measured activity-related features, knocking out Ubc7 in NPFR neurons led to a pronounced increase in the amount of time flies spent walking and performing turns, and to an overall increase in their average velocity compared to genetic controls (Figure 6F). Moreover, Ubc7 KO male flies exhibited increased social interactions between males, as shown by the higher levels of chase and song and reduced social clustering (Figure 6F). This suggests that protein ubiquitination in NPFR neurons is important for regulating the intensity of male-female and male-male sexual and social behaviors, and that Ubc7 is necessary to reduce male social interactions.

**Fig 6:**
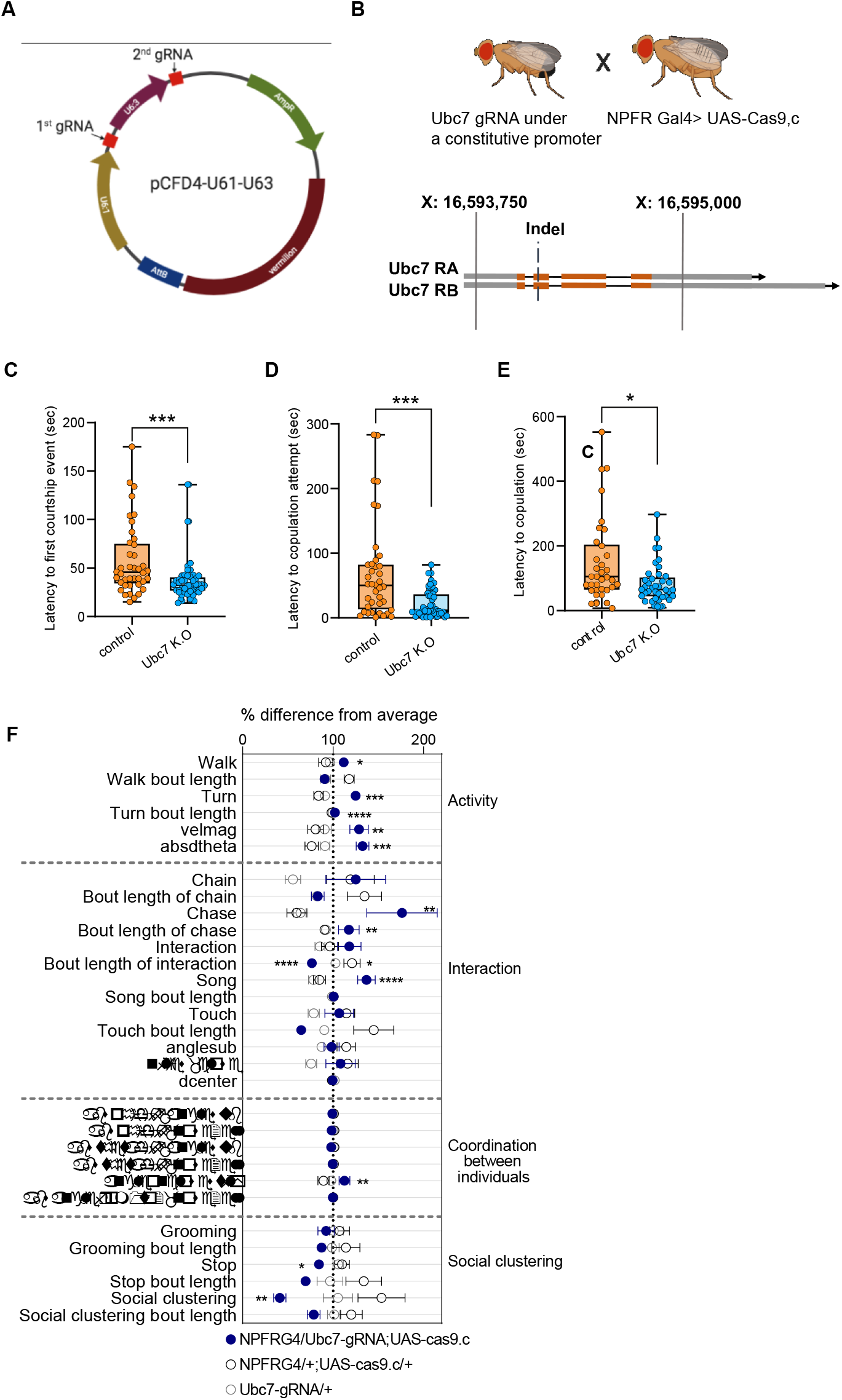
Tissue specific K.O of Ubc7 elevates the motivation to court and enhances male-male social interaction. **A** Representation of pCDF4 plasmid containing two gRNAs (red). **B** crossing scheme of flies containing gRNAs with NPFR-Gal4; Cas9.c flies to generate tissue specific Ubc7 K.O flies (upper panel). Lower panel depicts two Ubc7 isoforms (orange and grey blocks representing coding and non-coding exons respectively, Black lines representing introns). The double strand break occurred at the beginning of the 2nd coding exon. **C-E** Male flies containing NPFR-Gal4/Ubc7-gRNA; UAS-Cas9.c/+ (blue) were introduced to naïve females in courtship arenas and were video recorded, their courtship behavior was analyzed for latency to first courtship event (**C**), latency to first copulation attempt (**D**) and latency to copulation (**E**) compared to genetic controls (NPFRG4/attp1;UAScas9.c, orange). n=47 and 40 in C, n=46 and 39 in D, n=39 and 33 in E for experimental and control groups, respectively. Kruskal Wallis test with Friedman post hoc. *P<0.05, ***P<0.001. **F** Behavioral signatures of social group interaction in male flies harboring NPFR-Gal4/Ubc7-gRNA; UAS-Cas9.c/+ (blue) compared to NPFR-Gal4/+; UAS-Cas9.c/+ and Ubc7 gRNA/+ (grey and black, respectively). Data is presented as normalized scatter plots depicting % difference from average of 31 behavioral features. n=8, 5 and 11, respectively. One-way ANOVA with Tukey’s post hoc for normally distributed parameters and Kruskal-Wallis with Friedman test post hoc for non-normally distributed parameters. *P<0.05, **P<0.01, ***P<0.001, ****P<0.0001, n.s. P>0.05.

## Discussion

The intricate interplay between genes, neurons and behavior started to unravel decades ago with the Benzerian revolution in neurogenetics, and is still under intense investigation these days using plethora of tools in various model organisms. This study joins a growing avenue of studies that use contemporary genomic approaches to dissect the brain into units and illuminate their molecular content, as a step towards understanding the dynamic spatiotemporal environments in which genes function^15–17^. While many cell types exist in the fly brain, in this study, we analyzed only a small fraction of them, focusing mostly on NPFR neurons. Nevertheless, the transcriptome profiles of other neuronal populations in this dataset can serve as a resource for labs investigating other neurons.

We took two complementary pairwise based approaches to investigate the relative signature of NPFR: the first approach comparing profiles of each of the populations to those of all neurons and subsequently comparing the yes genes across populations; and the second approach performing pairwise comparisons between NPFR and each of the nine neuronal populations. Although the two approaches highlighted different number of differentially expressed genes, they resulted in similar hierarchical clustering patterns, and complemented the picture describing the distinct molecular landscape of NPFR neurons.

By comparing expression profiles of different neuronal populations to those of all neurons, we found that NPF expressing neurons represented a much more unique population than NPFR neurons. This may result from differences in cell number (40 NPF vs. ~100 NPFR cells) or may be associated with the heterogenous expression profile of NPFR as receptor neurons. The second explanation is supported by the enriched levels of receptors for neuropeptides and neuromodulators we detected, a finding that is in agreement with those of previous studies showing anatomical overlap between NPFR cells and some NPF and TH neurons^45,62^. At this point it is not known if NPFR neurons receive multiple inputs from many neuromodulatory systems, or whether they are composed of diverse groups of neurons with distinct combination of receptors. This question could be addressed in future studies by dissecting NPFR neuronal population into smaller subsets of cells using genetic intersection approaches or by single cell RNA-seq analysis.

The second part of this study investigated the functional relevance of the intricate transcriptome identified with our genomic approach. We discovered that global perturbation of RNA editing and protein ubiquitination programs in NPFR neurons resulted in dramatic behavioral phenotypes. Tissue-specific knockout of Ubc7 in male flies resulted in a strong motivation to court female flies, which contrasts with the complete loss of courtship behavior documented in male flies that lack Ubc7 in all cells^83^. This apparent discrepancy can be easily explained by distinct roles of Ubc7 in different tissues, the lack of which in NPFR possibly perturb the proper function of NPFR in restraining courtship as shown by Liu et al.^46^. Interestingly, a subset of NPFR-dopamine neurons has been shown to promote mating drive^45^, strengthening the notion that different sub-populations of NPFR neurons have distinct roles in regulating the motivation to court, and stressing the need for dissecting NPFR cells into smaller groups of neurons to analyze their transcriptomes and functions.

In addition to the increased motivation to court, we also documented increased frequencies of male-male social interactions, including increased levels of song and chase behaviors, which are normally absent in socially experienced male flies that are housed in groups^79^. This manifestation, together with the increased walking velocity and lack of social clustering behavior observed in groups of Ubc7 KO flies, resemble the behavioral properties of male flies exposed to social isolation, a condition that is known to promote aggression^84,85^. Given that the physical features of the FlyBowl set-up prevents the expression of aggression displays such as lunging, the increased chase behavior documented in the FlyBowl setup may be indicative of the male-male aggressive behavior that is normally suppressed by NPF action on NPFR neurons^86^ and that was promoted by perturbing the delicate proteome balance in NPFR neurons.

We have previously shown that different neuronal populations possess distinct repertoires of RNA editing levels in different transcripts^8^, suggesting that RNA editing may account for some functional differences between distinct populations in the brain. The pronounced behavioral outcome of perturbing RNA editing in NPFR neurons supports this hypothesis and shows that RNA editing is necessary for the proper function of these neurons. The phenotypic resemblance to Ubc7 K.O in NPFR neurons suggests that both manipulations perturb the function of NPFR in regulating social interaction, but closer inspection reveals the existence of interesting differences. While perturbing protein ubiquitination affects activity/arousal that may stimulate chase behavior, lack of RNA editing leads to pronounced increase in approach behavior, interaction, chase and chaining behaviors without changing activity levels. These differences suggest that perturbing RNA-editing and protein ubiquitination do not lead to global malfunction of NPFR but rather affect distinct targets that regulate different features of NPFR physiology and function.

The behavioral phenotypes of reducing ADAR levels were more pronounced in NPFR than in CRZ neurons, suggesting that RNA editing does not have a uniform role in all neurons but rather shapes the proteomic repertoire of different neurons to allow their distinct function. Our findings join a previous study that demonstrated the spatial requirements of ADAR expression in regulating locomotor behavior^72^, emphasizing the need to extend this to other behavioral paradigms, neuronal populations and even to studying the tissue specific role of specific editing events.

To conclude, in this study we demonstrated that the function of NPFR neurons depend strongly on the integrity of its transcriptome and proteome and is required to suppress certain neuronal programs that execute social behaviors. It will be interesting to explore whether this is also applies to other motivational and reward related behaviors based on the well-established role of NPF/R systems in processing of natural and drug rewards. It would also be interesting to use the INTACT methodology to examine dynamic changes in NPFR expression pattern under various conditions corresponding to different internal states.

## Methods

### Fly lines and culture

*Drosophila melanogaster* CS flies were kept in 25C°, ~50% humidity, light/dark of 12:12 hours, and maintained on cornmeal, yeast, Molasses, and agar medium. NPFR-GAL4 was a gift from the Truman lab (HHMI Janelia Campus), CRZ-GAL4 was a gift from the Heberlein lab (HHMI Janelia Campus), UAS-dicer, UAS-dADAR RNAi was a gift from the Lee lab (Stanford University), UAS-CAS9.c was a gift from the Schuldiner lab (Weizmann Institute). Vasa-CAS9 was a gift from Gershon lab (Bar-Ilan University), y^1^w^67c23^;P{CaryP}attP1 (BestGene BL#8621).

### Determining gene expression levels from RNA-seq

Previously published RNA-seq data was used^8^. Reads were trimmed using cutadapt^87^ and mapped to *Drosophila melanogaster* (BDGP6) genome using STAR^88^ v2.4.2a (with EndToEnd option and outFilterMismatchNoverLmax was set to 0.04). Counting proceeded over genes annotated in Ensembl release 31, using htseq-count^89^ (intersection-strict mode). DESeq2^90^ was used to measure differential expression analysis with the betaPrior, cooksCutoff and independentFiltering parameters set to False. Genes were filtered in a pairwise manner according to the following parameters: log_2_-fold change of at least |1|, adjusted P value lowers than 0.05 (Benjamini and Hochberg procedure) and a minimum of least of 30 normalized counts in one of the repeats.

### FlyBowl

FlyBowl experiments were conducted as described in Bentzur et. al^79^. In brief: groups of 10 male flies, which were socially raised in groups of 10 for 3-4 days, were placed in FlyBowl arenas, and their behavior was recorded at 30 fps for 15 minutes and were tracked using Ctrax^91^. Automatic behavior classifiers and Per-frame features were computed by JABBA^92^ tracking system. Data of all behavioral features was normalized to % difference from the average of each experiment for visualization.

### Courtship assay (ubc7)

4-5 days old naive males were placed with 4-5 days old virgin females in round courtship arenas (0.04 cm3 in volume), one male and one female in each arena. Courtship arenas were placed in behavior chambers, under controlled temperature and humidity (25°C, 70% humidity). Behavior was recorded for 10 minutes from the introduction of male and female pairs using Point-Grey Flea3 cameras (1080×720 pixels at 60 fps). Latency to copulation attempt and latency to copulation were quantified for each pair relative to the first wing vibration the male exhibited. Statistics: Kruskal Wallis test with Friedman post hoc.

### Generation of gRNA, transgenic constructs and transgenic flies

gRNA sequences were selected using the Fly-CRISPR algorithm (http://flycrispr.molbio.wisc.edu/), contain 20 nucleotides each (PAM excluded), and are predicted to have zero off-targets. Two different gRNA sequences were selected for Ubc7, both within the coding region of the gene, but not overlapping each other. Both gRNA sequences were cloned into the pCFD4 plasmid (Figure 6A). Cloning into pCFD4 was done using Q5^®^ High-Fidelity DNA Polymerase (BioLabs). gRNA-harboring constructs were injected to Drosophila embryos and integrated into attP landing sites using the φC31 system into attP1 (BL#8621) on the second chromosome. Injections were performed as services by BestGene (https://www.thebestgene.com/).

#### gRNA sequences

GTTAACACTTGACCCGCCCG

GCCCCATCAGCGAGGACAAC

### Generation of the germline ubc7 indel mutant

Transgenic flies expressing gRNA pCFD4 were crossed to flies expressing Vas-Cas9. Flies containing both the gRNAs and nos-Cas9 were crossed to a Fm7a balancer line, offspring were then collected and checked for the presence of an indel using DNA seq. The resulting indel is a deletion of 18-23 (Figure 6B, S3).

#### Primers foe DNA sequencing

Forward: AGAAAGCCACTCGATTCATTCGATA

Revers: GTCCAGAGCGTGGAGAAGAT

### Generation of tissue specific CRISPR

Transgenic flies expressing gRNA pCFD4 were crossed to flies expressing NPFRG4/+;UAS-Cas9.c/+.

### Statistical analysis

Data of each behavioral feature per experiment was tested for normality and consequently tested by either One-way ANOVA or Kruskal-Wallis tests followed by Turkey’s or Friedman post hoc tests using Prism. Statistical overrepresentation was generated usingPANTHER^93,94^ (http://pantherdb.org/citePanther.jsp)

### Graphics

Figure 6A, B were Created in BioRender.com

## Acknowledgments

We thank all members of the Shohat-Ophir lab for fruitful discussions and technical support. We express special thanks to C. Andrew Frank (University of Iowa) and Dion Dickman (University of Southern California) for their productive suggestions. This work was supported by the Israel Science Foundation Grant 384/14 and Israel Science Foundation Grant 174/19.

---

**Figure S1.**
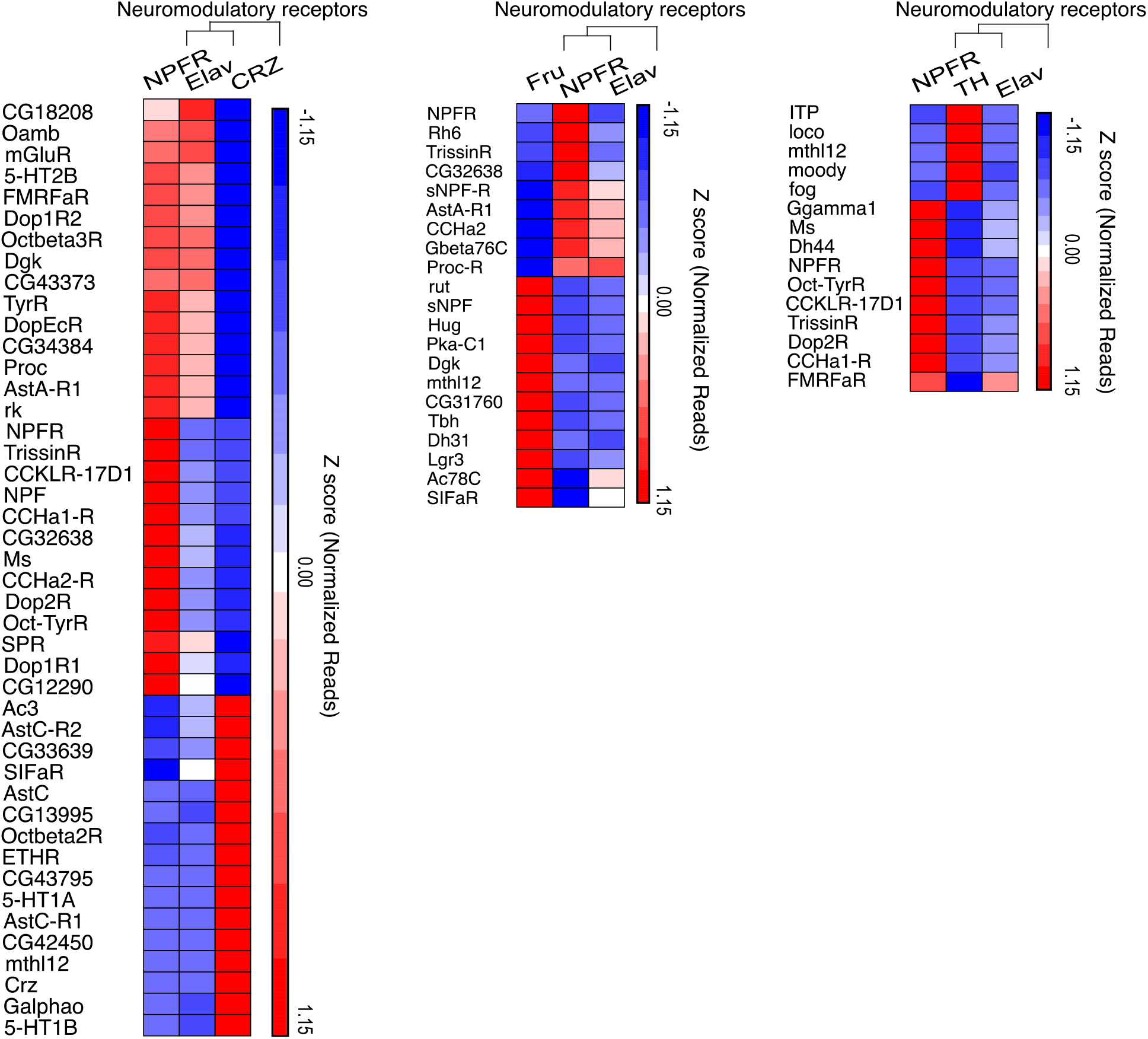
NPFR expressing neurons exhibit intricate expression patterns of receptors for neuropeptides and neuromodulators.

**Figure S2.**
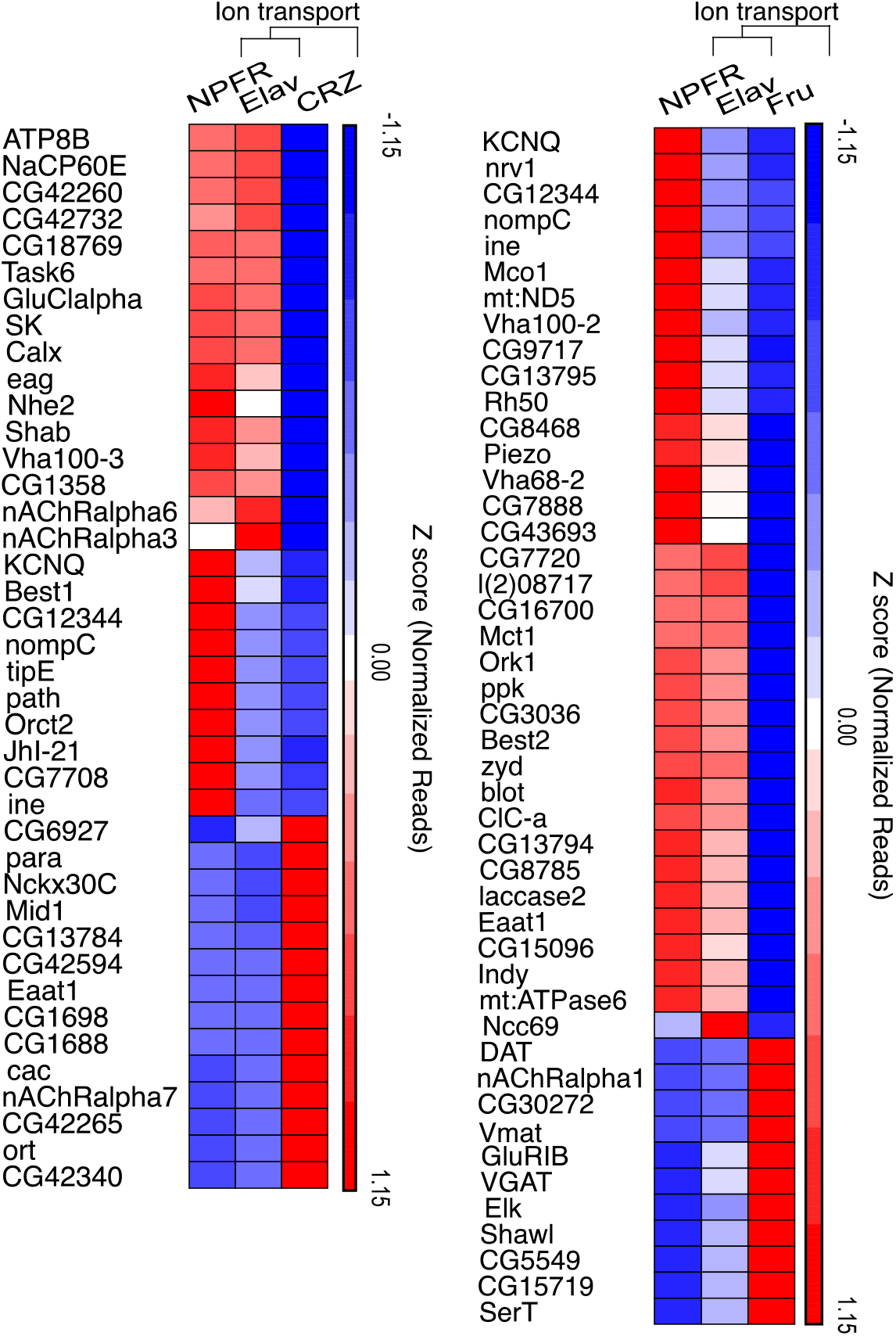
NPFR expressing neurons exhibit intricate expression of ion channels.

**Figure S3:**
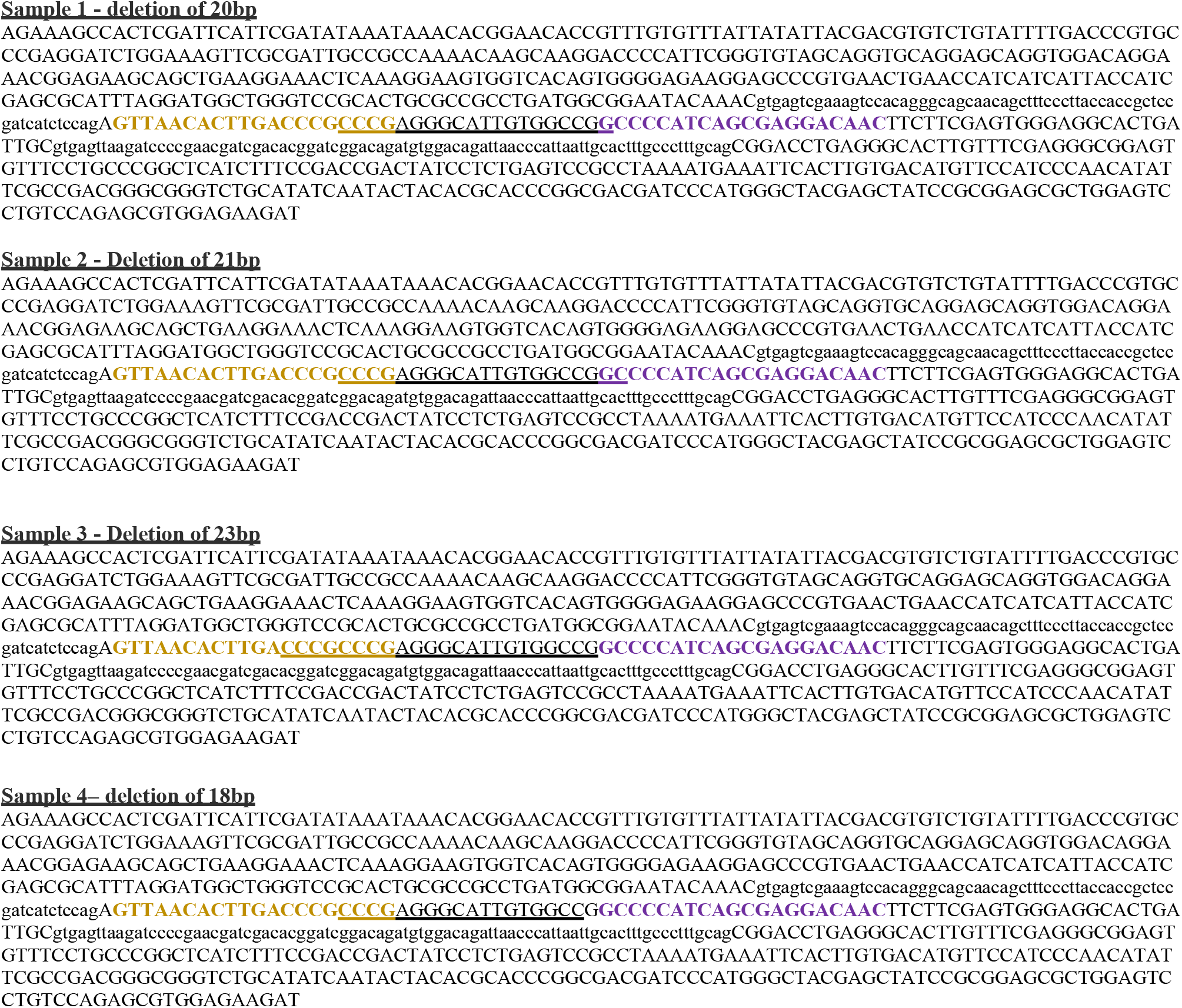
Sequenced 750 bp of Ubc7 DNA displaying 18-23 bp deletion in 4 male flies harboring Vas-Cas9; Ubc7 gRNA. Yellow and purple colors represent gRNA1 and 2 complementary sequences. Underlined sequences represent the deleted sequences.

